# An anti-depressant drug vortioxetine suppresses malignant glioblastoma cell growth

**DOI:** 10.1101/2024.03.04.583265

**Authors:** Mayur Nimbadas Devare, Matt Kaeberlein

## Abstract

Glioblastoma (GBM) stands as the predominant primary malignant brain tumor in adults, characterized by an exceedingly grim prognosis. Urgent efforts are essential to pioneer effective therapeutics capable of addressing both the intrinsic and acquired resistance exhibited by GBM towards existing treatments. This study employs a drug repurposing strategy to explore the anti-cancer potential of vortioxetine in malignant U251 and T98G glioblastoma cells. Findings from WST-8 cell counting assay and clonogenic assays indicated that vortioxetine effectively suppressed the short-term viability and long-term survival of glioblastoma cells. We also showed that vortioxetine inhibited migration of glioblastoma cells as compared to the control. Our findings encourage further exploration and validation of the use of vortioxetine in the treatment of glioblastoma.

## Description

Glioblastoma multiforme (GBM) stands out as the prevalent and most aggressive primary brain tumor among adults (Hanif, Muzaffar, Perveen, Malhi, & Simjee Sh, 2017). Despite extensive research and numerous clinical trials over decades (Alomari et al., 2021), the median survival rate remains stagnant at around 14 months. This challenge is compounded by the highly invasive nature of GBM cells, complicating efforts for complete surgical removal. Moreover, GBM cells demonstrate a propensity for developing resistance against the current multimodal treatment approach, incorporating the alkylating agent temozolomide (TMZ) and radiation therapy (Tan et al., 2018). This underscores the urgent need for novel treatments. Nonetheless, the extensive duration needed for the development of new small molecules and their validation for efficacy and safety in preclinical models presents a significant barrier to discovering novel therapies for this debilitating condition.

Drug repositioning may be one means of expediting therapeutic drug development for GBM. Drug repurposing involves utilizing previously approved drugs for new indications beyond their original use. There has been an increasing interest in drug repurposing across various medical domains (Hernandez et al., 2017; Nowak-Sliwinska, Scapozza, & Ruiz i Altaba, 2019). Repurposing drugs offers several advantages over novel drug discovery. Given that these drugs have already undergone thorough investigation for safety profiles and pharmacokinetic properties (Bahmad et al., 2020; Fisher & Adamson, 2021), drug repurposing can prove to be considerably more cost-effective and less time-consuming than the process of discovering entirely new drugs (Tan et al., 2018).

Vortioxetine (Figure 1A) is a multifaceted antidepressant that targets both serotonin receptors and transporters, employing a dual pharmacological approach (Chen, Hojer, Areberg, & Nomikos, 2018) however it’s mechanism of action is not precisely known (D’Agostino, English, & Rey, 2015). A recent study has shown that antidepressants can manifest antitumor effects (Zhang et al., 2013). To determine the anti-glioma activity of vortioxetine in GBM cells, U251 and T98 cells were incubated with increasing vortioxetine concentrations for 72 h (Figure 1B). All the cell lines were sensitive to treatment with vortioxetine in a dose-dependent manner. Using the cell counting kit-8 assay, the IC50 value of reduced viability for vortioxetine was 9 μM in U251 cells, 11 μM in T98 cells (Figure 1C,D).

**Figure 1.**
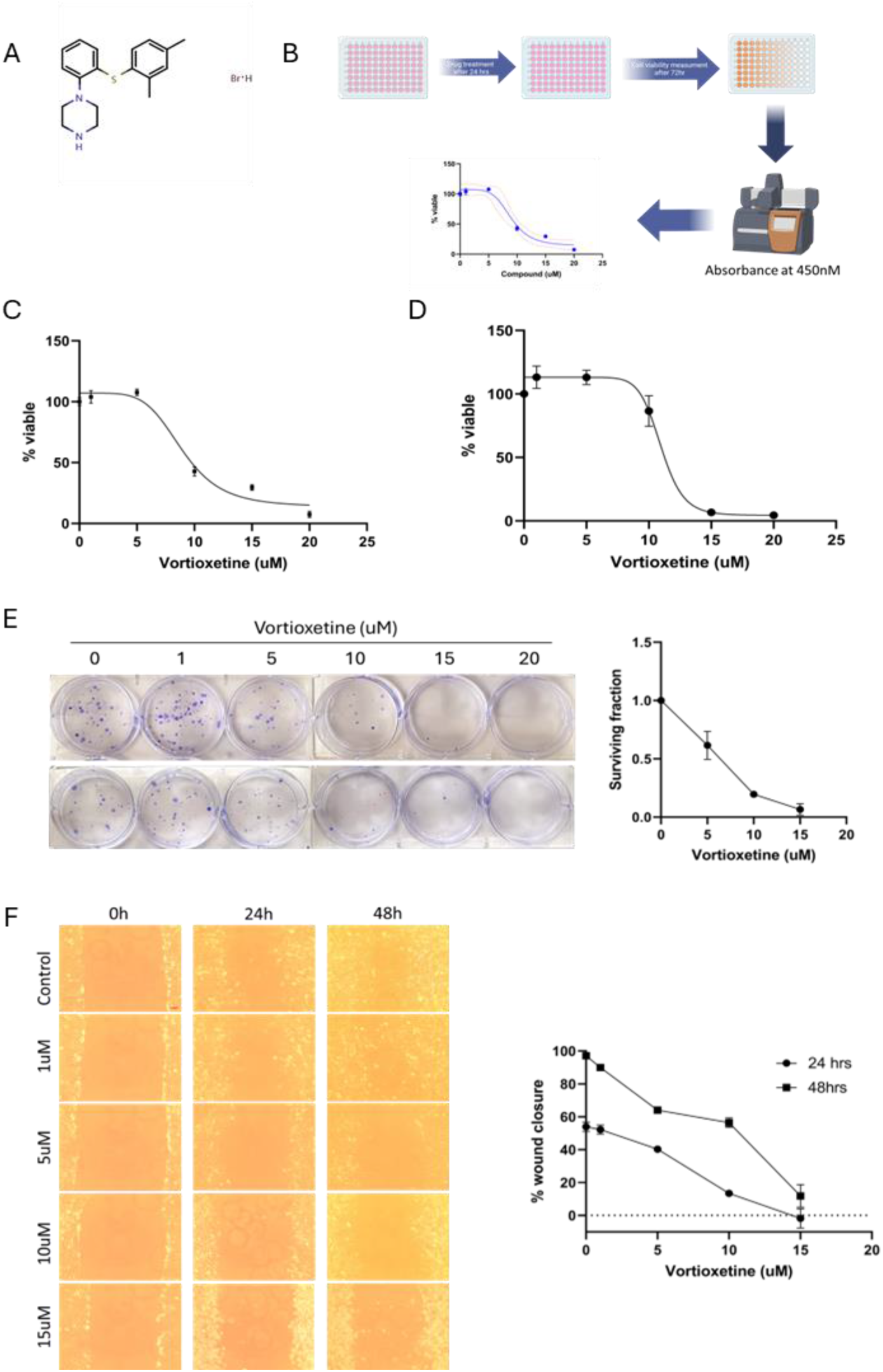
Vortioxetine suppressed glioblastoma cell growth. **(A)** Molecular structure of Vortioxetine **(B)** Depiction of screening for the effect of drug on glioblastoma Cell line. Cell viability was accessed by CCK-8 in U251 **(C)** and T98G **(D)** glioma cells. **(E)** Clonogenic assay was performed in U251 cells. After the experiment cells were stained with crystal violet, dried, and photographed. **(F)** The influence of vortioxetine on the migratory ability of glioblastoma cells at 24 and 48 h. The values shown are the means and standard deviation from three independent experiments.

The clonogenic assay was employed to assess the colony-forming capability of glioblastoma cancer cells. Following exposure to varying doses of vortioxetine (ranging from 1 to 20 μM), cells were incubated for 3 weeks. The findings revealed a notable reduction in clonogenic potential upon vortioxetine application, as indicated by diminished blue staining observed in the 6-well plates (Figure 1E). Therefore, along with proliferation rate of cell the clonogenic potential was also reduced to minimum by vortioxetine.

Cell migration plays a crucial role in the distant metastasis of cancer cells. To study the effect of vortioxetine on cell migration, it was evaluated using a wound healing assay. The results of the wound healing assay demonstrated a significant reduction in cell migration upon treatment with vortioxetine in U251 glioma cells. While the wound width in control cells was nearly closed, the wound width in treated cells remained unchanged at higher doses of the molecule (5-15 μM) (Figure 1F). Thus, the wound healing assay provided evidence that vortioxetine effectively minimized the migratory potential of the cells.

Overall, vortioxetine demonstrated significant suppression of both the short-term viability and long-term survival of glioblastoma cells. As compared to vortioxetine (Figure 1C, D), the IC50 value of TMZ, the gold standard therapy for GBM, is more than 1000uM in U251 and T98 cells (Natsume et al., 2005). This shows that vortioxetine is much more potent than TMZ in these cells. Additionally, our study revealed that Vortioxetine effectively inhibited the migration of glioblastoma cells compared to the control group. These results advocate for continued investigation and validation of vortioxetine’s potential application in glioblastoma treatment.

A few anti-depressants have shown to possess anti-tumor activity through various mechanisms. These mechanisms include actions involving mitochondria (Yuan et al., 2015) and Ca2+-mediated cell apoptosis (Zhang et al., 2013), nerve growth factor receptor (NGF receptor)-mediated, and Fas death receptor-mediated cell apoptosis (Yuan et al., 2015). Additionally, antidepressants impact pathways associated with JNK/c-Jun, PI3K/AKT/mTOR, NF-kappa B, and ERK signaling, as well as exert antiproliferative effects via modulation of cell cycle regulators. Moreover, antidepressants are observed to inhibit tumors by altering the immune response or microenvironment of cancer cells (Zhang et al., 2013). Recent studies have shown that vortioxetine suppresses gastric cancer cell growth via JAK2 and SRC proteins (Li et al., 2023) and promotes apoptosis and autophagy through PI3K/AKT pathway (Lv et al., 2020). Further experimentation is warranted to elucidate the molecular mechanism by which vortioxetine suppresses glioblastoma cell growth.

## Materials and methods

### Cell lines and culture conditions

The human malignant glioblastoma cell lines U251 and T98G were kindly provided by Dr. Nephi Stella, Departments of Pharmacology, University of Washington. Cells were grown in a monolayer using Dulbecco’s Modified Eagles Medium (DMEM) supplemented with 10% fetal bovine serum (FBS) (Gibco, Life Technologies Corporation, Saint Louis, USA) and penicillin-streptomycin solution (Gibco, Life Technologies Corporation, Saint Louis, USA). Cells were maintained at 37 °C, in a humidified atmosphere consisting of 5% CO_2_ and 95% air. Cell growth medium was changed routinely every two to three days and sub-culturing of cells was performed with a solution of 0.25% trypsin EDTA (Sigma -Aldrich, Saint Louis, USA).

### Treatment

The quantity of vortioxetine (Cat. No. P50895, Achemblocks, CA, USA) required to make a stock solution of 10 mM was weighed and dissolved in appropriate volumes of Dimethyl sulfoxide (DMSO) (Sigma-Aldrich, USA). Aliquots of 10 μL of the compound were stored in Eppendorf tubes at − 20 °C for use within one-month to prevent repeated freeze and thaw cycles.

### Cell viability assay

The WST-8 cell counting kit (ApeXBio, USA) was used to determine cell viability according to manufacturer’s protocol. Cells were seeded in sterile 96-well cell culture plates at a density of 5000 cells per well and allowed to settle overnight after which they were exposed to increasing concentrations of vortioxetine (0 μM to 20 μM) for 72 h, with 0 μM serving as vehicle control (equivalent amount of DMSO in the 20 μM concentration). Cells were incubated with 10 μL of the CCK-8 solution for 1 h 30 min at 37 °C and mixed gently on an orbital shaker for homogenization. Measure the absorbance at 450 nm using a microplate reader (BioTek Epoch Microplate reader, Agilent tech) and the mean cell viability was calculated relative to control. The maximal inhibitory concentration value (IC50) was determined through a survival curve using GraphPad Prism 9 software (GraphPad software, San Diego, CA, USA) from three experimental repeats performed in triplicate wells.

### Clonogenic assay

2×10^4 cells in 2ml media were seeded in 6 well plates. After 24hrs, vehicle (DMSO) or vortioxetine (1,5,10,15,20uM) were added for 24hrs. After 24hrs media was collected in individual 15ml tube, attached cells were washed wish PBS and PBS was collected. Cells were treated with 300ul trypsin for 4-5 mins and 1ml media was added, cells were rinsed with media to get single cell suspension. All the cells were collected in respective 15 ml tubes. Centrifuge for 7 min at 200g and remove supernatant and resuspend cells in 1ml new media. Take 6ul from each tube and count the number of cells by hemocytometer. Re seed in three different 6 well plates (triplicate) 200cells/well. Incubate the plates at 37C for 12 days. Media was removed, washed with 2 ml PBS. Remove PBS and add 2ml of 6% glutaraldehyde, 0.5% Crystal violet in H2O. Incubate at RT for at least 30 min. Remove staining solution. Take tap water in ice bucket and submerge plates to wash off excess stain. Dry at RT and plates were photographed.

### Migration assay

10^5 cells were seeded in 2ml media were seeded in 6 well plates. After 24hrs, confluent cell surface was scratched with the help of sterile 200ul pipette tip. Media was removed and cells were washed with PBS. Media with vehicle (DMSO) or vortioxetine (1,5,10,15uM) were added and 0h images were taken, and plates were incubated at 37C for 48h. Plates were removed from incubator after every 24h and images were taken on Amscope microscope. Distance between two ends of the scratch was measured with ImageJ software. Rate of cell migration = (initial wound width-final wound width)/time. The graph was plotted using GraphPad Prism 9 software (GraphPad software, San Diego, CA, USA) from three experimental repeats performed in triplicate wells.

